# Real-time ultrafast holography of focused ultrasound wave fields using an ultrasound scanner

**DOI:** 10.64898/2026.10.15.759327

**Authors:** Kazuhiro Matsui, Françoise Chavrier, Takashi Azuma, Ichiro Sakuma, W. Apoutou N’Djin, Rémi Souchon

## Abstract

Acoustic pressure field measurements are required for marketing clearance of medical ultrasound instruments, but the gold standard method, hydrophone scanning, is time consuming.

The objective of this work was to develop an ultrafast holographic method for real-time imaging of focused ultrasound fields. The technique is based on parallel acquisition with a sensor array, and FUS field reconstruction based on time reversal assuming a homogeneous lossless medium. The geometric characteristics of the sensor to enable accurate reconstruction were evaluated through numerical modelling. A practical example involving a commercially-available 128-element linear-array ultrasound imaging probe is presented. Finally, preliminary experiments were conducted on in-vitro soft tissues to validate the concept of simultaneous dual-mode imaging, combining conventional B-mode tissue imaging with FUS field holography.

In modelling of ideal sensor characteristics (2D matrix; intercepted energy > 99%), parallel holography allowed reconstructing wave fields with excellent agreement with the Rayleigh Integral reference (errors << 0.01%). Successfully experimental demonstration of real-time ultrafast wave field holography was achieved with the commercial imaging probe (7-10% error in the -6dB focus vs hydrophone measurements, acquisition time: 100 µs, display rate: 15 frames/s), despite a suboptimal intercepted energy (44%). Real-time dual-mode imaging of continuous wave fields allowed targeting localized tissues regions, while holographic movies highlighted the propagation of transient waves.

Overall, real-time ultrafast holography offers a practical way to perform metrology and guidance of ultrasonic devices, by visualising waves fields typically used for ultrasound imaging, or those used during thermal or non-thermal ultrasound therapy.

A CC-BY public copyright license has been applied by the authors to the present document and will be applied to all subsequent versions up to the Author Accepted Manuscript arising from this submission, in accordance with the grant’s open access conditions.

## I. Introduction

Diagnostic and therapeutic ultrasound utilize ultrasound fields whose shapes, amplitudes and timings are carefully designed to achieve the desired bio-effect. For example, appropriate spatial-temporal focusing of ultrasound fields enables high resolution ultrasound imaging [1] and targeted therapies [2], [3], [4]. The ability to visualize and quantify the ultrasound field is thus critical in order to ensure that imaging and therapy are safe, accurate and efficient.

To date, no method can measure the ultrasound fields in vivo. Moreover, simulation models fail to provide fully accurate predictions of these wave fields inside biological tissues, due to heterogeneity of the tissues, inter-individual variations, and uncertainties in the determination of their acoustic properties (e.g., attenuation, speed of sound). Additional aberrations may also occur because of partial reflection and refraction at tissue layers (e.g., muscle, skin or fat), of anatomical barriers (e.g., bones or gas), and eventually of imperfect acoustic coupling between the ultrasound source and the tissues.

In homogeneous fluids however, the ultrasound field can be predicted accurately by theoretical models [5], [6], and measured experimentally using hydrophones [7], [8], [9]. However, hydrophone measurements require point-by-point scanning. Scanning a 3D volume may take hours, even days, depending on the field of view (FOV) and sampling interval.

Acoustic holography [10], [11], [12], [13], [14], [15] is also based on hydrophone measurements. It allows 3D reconstruction while scanning only a single 2D plane. However, it remains time consuming (several hours) because it requires large FOV and dense sampling (typically half a wavelength or less).

Optical methods [16], [17], [18], [19], [20], [21], [22], [23] can measure a 2D projection of the instantaneous ultrasound field in real time, but only in optically transparent media, thus not in biological tissues. They are based on the local modification of the optical refraction index, induced by the pressure wave. Three-dimensional ultrasound fields can be obtained by the use of computed tomography algorithms, but by losing the real-time advantage.

Motion-sensitive magnetic resonance imaging pulse sequences can measure the ultrasound pressure field inside biological tissues [24], [25], [26]. However, the technique cannot be used in vivo because gradient amplitude and switching rate exceed safety limits.

Backscattered ultrasound imaging [27], [28], [29], [30] can image the ultrasound field using ultrasound scanner. However, the technique is not quantitative, because the pressure field images are biased by local variations in the backscattering coefficient of the tissues.

This study introduces an ultrafast holographic method for imaging focused ultrasound (FUS) wave fields in real-time. It is based on acoustic holography, replacing the hydrophone by a multi-element ultrasound probe driven by an ultrasound scanner, and positioning the sensor so that most of the FUS energy flux is intercepted by that sensor.

## II. Principles and theory

### A. Kirchhoff integral, reciprocity principle, time reversal

Acoustic holography is not a new concept. It has been described and implemented on numerous occasions, with variations in the assumptions, in the numerical implementations (in the time domain or in the frequency domain), in the accuracy of the results, and in the level of details in the descriptions of the methods. The implementation that was used in the present study is based on the work by Sapozhnikov et al. [13], and provides the exact solution in a lossless homogeneous fluid.

A wave field can be reconstructed from point measurements using the Kirchhoff integral theorem [31], [32]. The theorem states that the velocity potentials of the wave field can be calculated at any observation point by integrating the individual contributions of each elementary area of any arbitrary surface that encloses the observation point (Fig. 1a). The surface needs to be a closed surface, so that the entire acoustic energy flux passing through the surface is known. The inner medium is assumed to be lossless, homogeneous, linear, and isotropic. This theorem is the basis for the Rayleigh integral, wherein the closed surface embeds the acoustic source [33], [34].

**Fig. 1.**
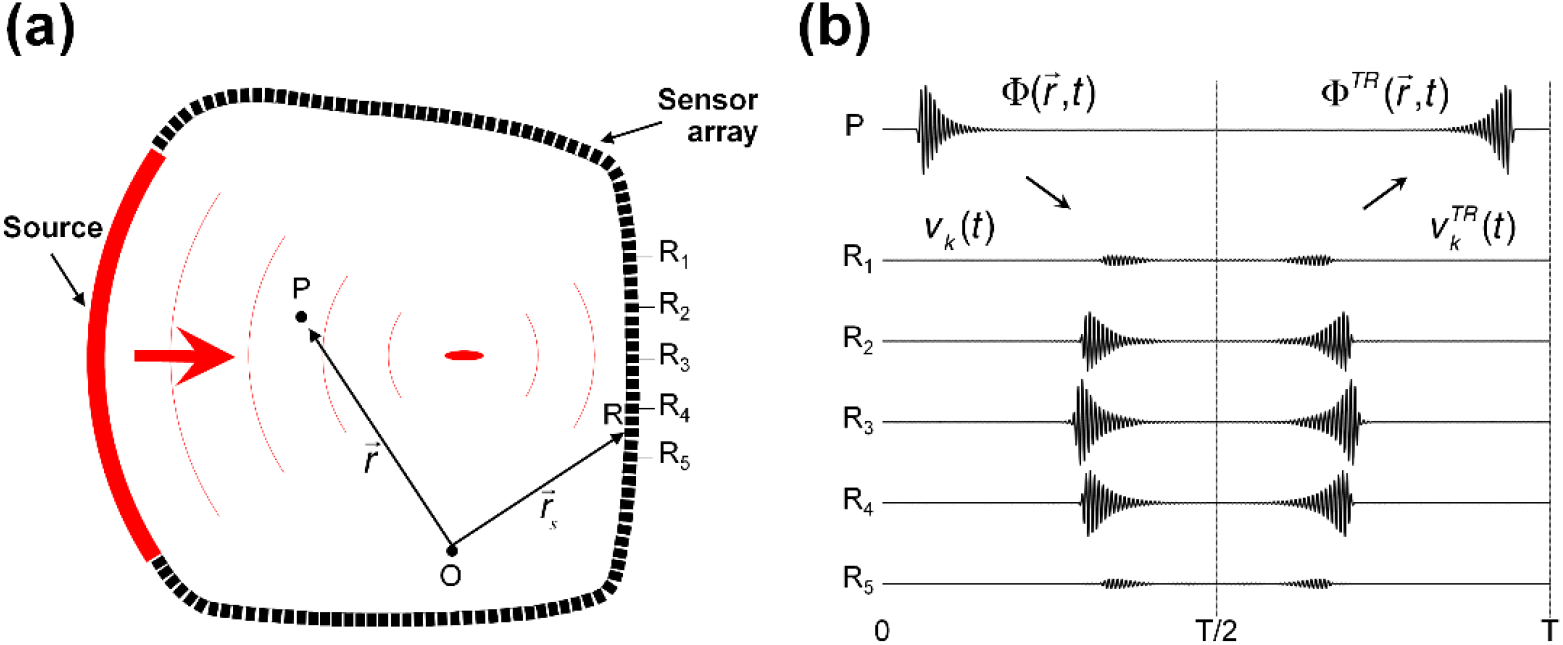
Conceptual schematics of the method for imaging ultrasonic fields based on a time reversal principle. **(a)** Imaging system configuration. An ultrasound source generates a field, and a sensor array (R_1_, R_2_, …, R_N_) placed on a closed surface captures the incident field. The point O is the origin of the coordinate system. An arbitrary observation position 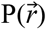 and sensor array positions 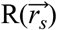 are shown. **(b)** Timing sequence. During the first part of the experiment (0 ≤ *t* ≤ *T*/2), the sensor array is used as a receiver. The incident wave propagates in the medium, and the observation point P undergoes temporal variations in velocity potential 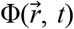 (shown) – note that corresponding changes in pressure and in particle velocity exist, but are not shown here. The normal velocity waveforms – *v*_*k*_(*t*) – undergone by the sensor elements are recorded. In the second part of the experiment (*T*/2 ≤ *t* ≤ *T*), the sensor array is used as a source. Time-reversed normal velocity waveforms 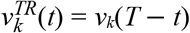 are re-emitted back into the medium. This time-reversal process generates a new “time-reversed” acoustic field 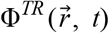, propagating backward toward the source, which follows 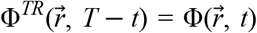. In other words, the observation point P undergoes the original vibration pattern again, except that it is now played backward in time.

The reciprocity principle states that, in an acoustic experiment, the role of an emitting surface (i.e., source) and of a receiving surface can be interchanged [35]. In the Kirchhoff integral theorem, the role of the surface is irrelevant; i.e., the surface can be a source, or a sensor, or just any point in space. Hence, if the surface is a set of point-like sensors, say a sensor array, the theorem still applies.

The combination of reciprocity and of the Kirchhoff integral theorem is the basis of Time Reversal (TR). Time Reversal was initially applied for generating focused ultrasound fields in complex media [36], [37], and in acoustic holography. Time Reversal was also used for imaging wave fields or sources in acoustics [38], [39], [40], [41], in seismology [42], [43], [44], in electromagnetics [45], [46], and in optics [47], [48].

In subsequent paragraphs, we will denote “Time Reversal experiment” whenever the wave is actually generated physically, and “calculated Time Reversal” when the wave is not generated, but predicted.

### B. The Time Reversal experiment

In a time reversal experiment (Figure 1), point-like acoustic sensors (R1, R2, …, RN) are positioned on the closed surface. They measure the incident field 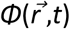 [36]. Each sensor converts the local pressure to an electrical voltage. The voltage waveforms *u*_*k*_(*t*) are recorded simultaneously on all sensor elements. Assuming that the response of the sensor elements is linear, the component of the particle velocity vector normal to the *k*^th^ sensor element, 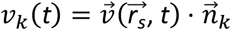, can be derived from the received electrical voltage *u*_*k*_(*t*):

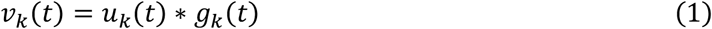

In subsequent paragraph, for the sake of simplification, the normal component of velocity is called “normal velocity”. A list of symbols is provided in Table I. Briefly, the symbol * represents the convolution operator, and *g*_*k*_(*t*) is the acousto-electrical impulse response, representing the conversion from mechanical energy to electrical energy of the *k*^th^ sensor element. Accurate measurement of the normal velocity *v*_*k*_ requires that each sensor element is at normal incidence relative to direction of propagation of the FUS beam, and small in size compared to the wavelength.

The signal acquisition is stopped at time T/2 (Figure 1b). Then, from time T/2 onward, the sensor elements are used as a new source. It is possible because most acoustic sensors are reversible. They can also convert an electrical signal into a mechanical displacement. All elements are driven simultaneously with electrical voltage *e*_*k*_(*t*):

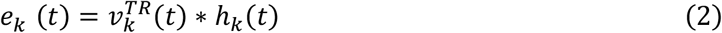

where *h*_*k*_(*t*) is the electro-acoustic response of the element (conversion from electrical energy to mechanical energy), and:

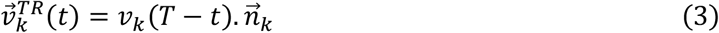

is the time-reversed normal velocity vector field to be generated at the surface of the *k*^th^ element. These waveforms are identical to the incident velocity waveforms *v*_*k*_(*t*), except that they are played backward in time. Note that for lossless sensors, such as reversible piezoelectric crystals would be, *g*_*k*_(*t*) * *h*_*k*_(*t*) = *δ*(*t*), and the driving voltage simplifies to *e*_*k*_(*t*) = *u*_*k*_(*T*-*t*). This process generates a new wave field *Φ*^*TR*^ in the medium. In a lossless medium, the new field is identical to the incident field, except that it is “time-reversed”, i.e. played backward in time [36], [37]:

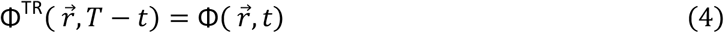

Time reversal therefore recreates the exact incident field, with the exception of evanescent waves. The latter can be ignored whenever the observation position is several wavelengths away from the new source [49].

### C. Calculation of the time-reversed ultrasound field

The time-reversed field *Φ*^*TR*^ can be created experimentally. Therefore, it can also be predicted, just like any wave field, using a model. Below, we will use the Rayleigh integral [33], [34], [50]. The sensor array is now the source, and the normal particle velocity vector field 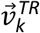 is applied at its surface. The velocity potential of the time-reversed field *Φ*^*TR*^ can be predicted at any observation position 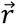 and any time *t* using:

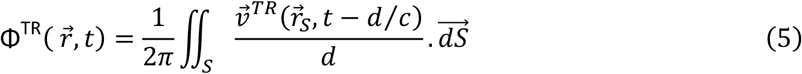

where 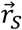 is the position of the elementary integration surface 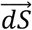, and 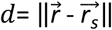 is the distance between that surface and the observation position 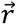. The integral is implemented numerically:

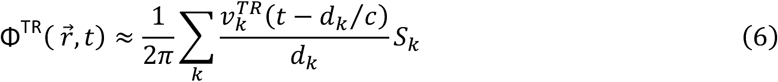

where the surface integral was replaced by a discrete sum over the elements of the sensor array, and *d*_*k*_ is the distance between the observation position 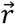 and the center of the *k*^th^ element. The approximation in equation (6) arises from the particle velocity being usually non uniform on the sensor element. According to Eq. (3), the time-reversed velocity waveforms 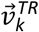 can now be replaced by the velocity waveforms *v*_*k*_ that were measured on the sensor elements:

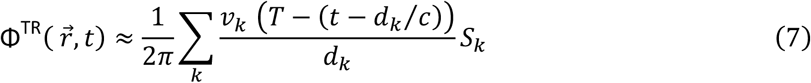

Finally, taking advantage of Equation (4), the time-reversed field *Φ*^*TR*^ can be replaced by the field of interest, 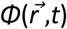:

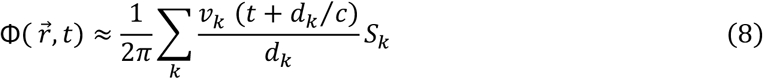

Equation (8) is a quantitative estimate of the velocity potential of the incident field. Other acoustic quantities such as pressure p, particle velocity 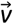 and all three components of the acoustic intensity (Poynting) vector 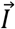 are easily derived [32]:

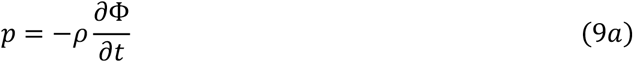

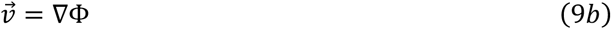

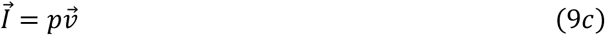

where *ρ* is the density of the medium. It is worth noting that equations (8) and (9) actually estimate the field at any observation position 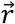 and any arbitrary time *t*. It is therefore possible to reconstruct a “holographic wave movie” showing the propagation of any transient or continuous wave in a region of interest, based on a single acquisition, then by varying time *t* in these equations. A prerequisite, however, is that velocity waveforms *v*_*k*_(*t*) have duration long enough to capture the entire wave, with no truncation in time.

### D. Summary of the underlying assumptions

The accuracy of the reconstruction depends on five hypotheses: (H1) The sensor array surface is large enough to captures the entire energy flux of the field; (H2) the dimension of the individual sensor elements is small compared to the wavelength, so that particle velocity can be assumed uniform on the element; (H3) the sensor elements are positioned normal to the direction of propagation of the FUS field, (H4) the sensor elements provide a linear response (see equation (1)), and (H5) the medium is homogeneous and lossless. It is thus interesting to observe that a FUS field in water can meet all these requirements, especially if the sensor is positioned in the vicinity of the focus, or even at the focus.

### E. Approximation using conventional beamforming

It is also interesting to note that equation (8) is similar to the dynamic receive focusing technique implemented in medical ultrasound imaging scanners. Ultrasound scanners create anatomical images by transmitting acoustic waves in the body, typically using a plurality of sensor elements. The raw echo signals *s*_*k*_(*t*) are recorded on each sensor element, and a beamformed image 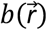 is generated by a delay-and-sum process called “dynamic receive focusing”, or “dynamic receive beam forming”, or conventional beamforming [51]:

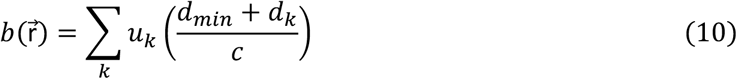

where *u*_*k*_(*t*) is the voltage signal measured on the k^th^ array element, *d*_*k*_ is the distance between the k^th^ array element and the observation position, and *d*_*min*_ = min(*d*_*k*_) is the shortest path between the array elements and the observation position. Note that *d*_*min*_ depends on the position, so it should be noted 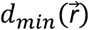. In the literature, the beamformed image 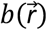 is often called the radio-frequency image or the RF image. The ultrasound image 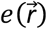 is the amplitude, or envelope, of the beamformed image.

A demonstration is given in Appendix A that the envelope of the wave field *Φ* can be estimated quantitatively using the ultrasound image:

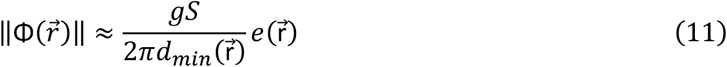

where *S* is the total area of the imaging probe, and *g* is the sensitivity of the elements (in units of V per m.s^-1^). Equation (11) is valid in the specific case of a continuous wave, or for burst waves with duration long enough so that the field is steady during the acquisition (i.e. for t∈[0,T/2]).

Appendix A also includes a method to remove the factor *d*_*min*_, using the appropriate time gain compensation (TGC) settings. As a consequence, the envelope of a continuous wave field can be observed directly and in real time using any conventional ultrasound scanner, taking advantage of the fast beamformers implemented in modern scanners. The accuracy of the wave field obtained with the conventional beamformer is not as good as that obtained with time reversal (Equation 8). The analysis in the Supplementary Materials shows that higher discrepancies are expected close to the imaging array.

### F. Dual-mode imaging (anatomy + wave field)

Some research-oriented scanners include a “passive imaging mode” wherein the transmission of ultrasound waves by the scanner can be disabled. These scanners operating in passive mode are fit for imaging the FUS wave field emitted by another transducer. In passive mode, the ultrasound imaging probe can be seen as an array of hydrophones for reconstructing in real time the FUS wave field. However, the vast majority of conventional medical ultrasound scanners are not passive receivers. They operate in active imaging mode, by transmitting ultrasound imaging waves into the body. These imaging wave will interact with the FUS wave field to be observed. The interaction between the FUS wave field and the imaging wave generated by the scanner needs to be studied and accounted for. At least two implementations are possible: interleaved dual-mode, and simultaneous dual-mode.

In interleaved dual mode, the scanner alternates between conventional active ultrasound imaging and passive wave field imaging. To do so, the transmission by the ultrasound scanner is disabled every second frame. Ideally the sequence is also synchronized with the FUS transducer, so that the FUS wave fields of interest is generated only when the scanner is in passive receiving mode. Two images are created: the anatomy image, using active imaging mode and conventional beamforming, and the FUS wave field image, using passive imaging mode and the Time Reversal equation (Equation 8). The two images can be displayed side by side, or they can be superimposed. The reconstructed wave field is exact, provided that all hypotheses H1-H5 are satisfied.

In simultaneous dual mode, the scanner receives the FUS wave field and the imaging wave field (the backscattered echo signals) simultaneously. When the FUS field has a narrow bandwidth, the two fields can be separated by filtering in the frequency domain, and each field can be processed separately, using the appropriate algorithm. Otherwise, the entire received signal is processed using conventional beamforming, resulting in an anatomic image with a superimposed FUS wave field (Equation 11). That reconstructed wave field is slightly inexact, but interesting because it is readily available on any commercially available ultrasound scanner.

## III. Materials and methods

Simulations were performed to demonstrate the feasibility and evaluation the accuracy of the ultrafast holographic technique when using an ideal ultrasound matrix probe (simulation A1), when using hypothetical suboptimal probes (simulations B1 and B2), and when using a commercially available linear probe (simulation B3). Then, experimental proofs of concept were performed using a linear array and a research-oriented ultrafast ultrasound scanner to demonstrate the feasibility of real-time ultrafast holography of FUS wave fields. All results were compared to hydrophone measurements, except for the transient wave movie that was only compared to simulations.

### A. Simulation of FUS field holography with an ideal imaging probe

Simulations A1 were performed in water, for both continuous wave (CW) and transient wave (TW) fields. The FUS source and the imaging sensor (receiver) array were positioned opposite to each other. In this first simulation, an ideal imaging sensor was simulated by an “optimal” 2D matrix whose characteristics are listed in Table II. The simulated configuration is shown in Figure 2. An orthonormal coordinate system is defined with x- and y-axis oriented respectively along the length and width of the imaging array, and z-axis oriented along the FUS propagation direction. The FUS source was a single-element spherical concave transducer with a centre frequency *f* = 2.4 MHz, a diameter of 70 mm, and a radius of curvature (focal length) *R*_*c*_ = 90 mm. The distance from the imaging sensor array to the focus of the source was *z*_*f*_ = 25 mm.

**Fig. 2.**
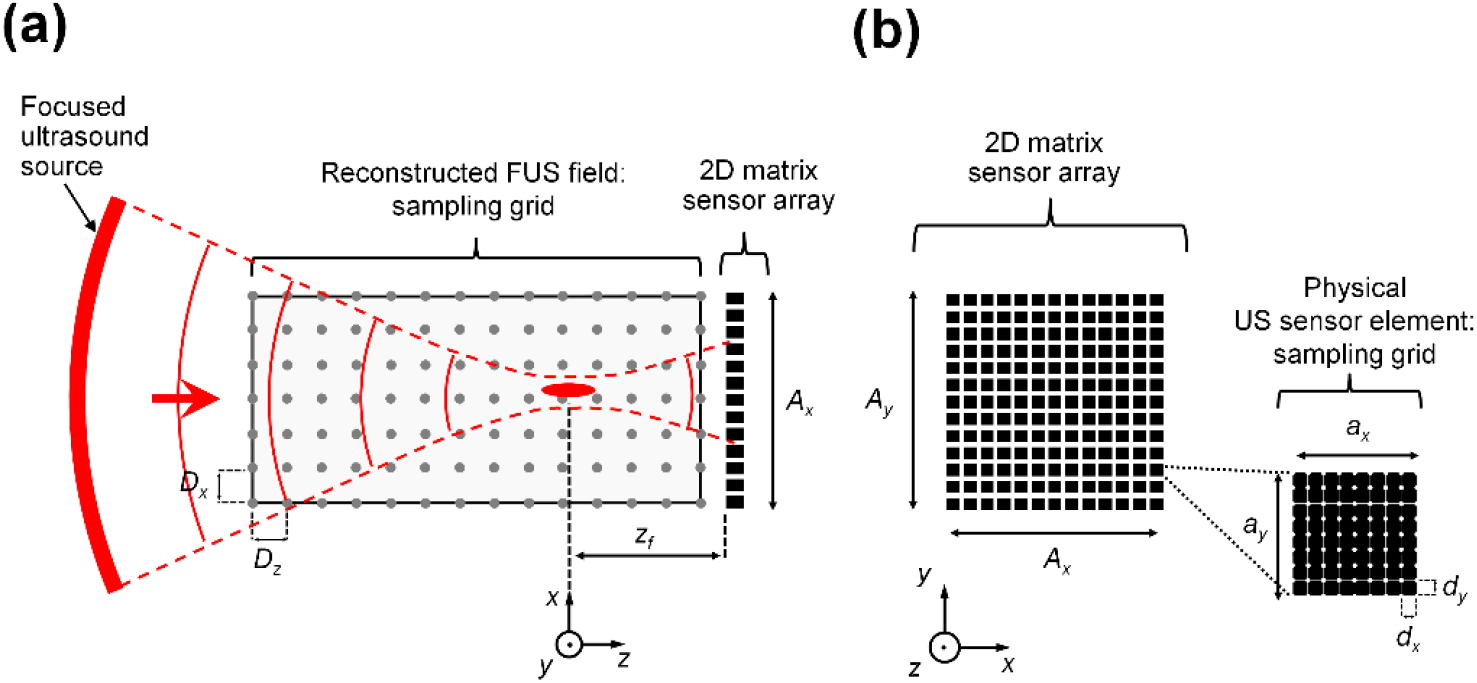
Simulation configuration. **(a)** A focused ultrasound source and a finite-aperture sensor array were positioned opposite to each other. Continuous and monochromatic waves generated from a focused ultrasound source were received by a 2D-element and square-shaped planar sensor array with a total dimension of *A*_*x*_ *× A*_*y*_ (nearly ideal conditions). Deviations to ideal conditions were progressively introduced until a suboptimal case of 1D linear imaging array. The distance from the sensor array to the focus of the source was *z = z*_*f*_ = 25 mm. 2D spatial sampling of the FUS field reconstruction: *D*_*x*_ *× D*_*z*_. **(b)** Each physical element of the imaging (sensor) array had a dimension of *a*_*x*_ *× a*_*y*_. 2D spatial sampling of an ultrasound element: *d*_*x*_ *× d*_*y*_. Numerical values are available in **Table II**.

#### 1. True wave field

The wave field of interest 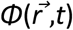 was calculated using the Rayleigh integral for continuous waves fields, and using the impulse response method for transient wave fields [52]. On the FUS source side, the normal velocity at the transducer surface was assumed to be a CW monochromatic 2.4 MHz sine wave. The acoustic power was set to 1 W. The medium was assumed to be lossless and homogeneous (water at 20°C, with speed of sound *c* = 1482 m/s [53]. The sampling frequency was set at 153.6 MHz (64 times the frequency of the source). The spatial sampling was dx = dy = 50 µm (corresponding to λ/12).

#### 2. Received voltage *v*_*k*_(*t*) on the sensor elements

For each element, the normal components (to the surface of the imaging element) of the particle velocity vectors of the incident FUS field, 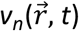, were calculated on all grid points located on the element. The eventual partial reflection of the incident FUS wave on the receiving array was ignored. Then, the normal velocity waveform *v*_*k*_(*t*) associated with the *k*^-th^ element was calculated by averaging 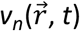 over the surface of the element. The sensitivity *g*_*k*_ was arbitrarily set to 1 m.s^-1^.V^-1^.

#### 3. Estimation of the wave field

The image of the wave field, 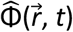, was created using the time reversal algorithm (Equation 8). The images obtained of the CW or the TW FUS fields were then directly compared to the “true fields” simulated in the first step.

### B. Simulation of FUS field holography using suboptimal probes

Ultrasound probes have a limited surface, or “footprint”. For example, the footprint of a commercially-available Verasonics / Vermon M3dV matrix (1024 elements) is only 1×1 cm. Hence the optimal condition of a sensor surface encompassing the entire volume of interest, as shown in Figure 1, cannot be satisfied. Moreover, the elements cannot be made infinitely small, and are often rectangular. As a consequence, errors are to be expected in the wave field reconstruction.

Nevertheless, conventional ultrasound imaging probes are worth investigating, because they are readily available. They can potentially allow FUS field imaging, provided that their limitations are known. This is why, in this second set of simulations, we investigated the loss in accuracy when either the total surface, or the element size, or both, are suboptimal. The details of the simulated sensors are given in Table II.

#### 1. Simulation B1. Effect of suboptimal sensor surface

In scenario B1, the dimension of the matrix was progressively reduced until only a small fraction of the total energy flux was captured. The elements are still smaller than a half-wavelength.

#### 2. Simulation B2. Effect of suboptimal element size

In scenario B2, the matrix was large enough to allow the total energy flux of the FUS field to be captured. However, the size of the square elements was progressively increased.

#### 3. Simulation B3. Using a commercially available linear array

Finally, in scenario C, the elements were rectangular. The pitch was smaller than half-wavelength, but the element height was much larger than the wavelength, resulting in simultaneous violation of requirements H1 & H2. This scenario corresponded to the specific case of a commercially-available 1D-linear imaging probe used in subsequent experiments. The probe exhibits ideal sensor length (38.4 mm) and element size (0.3 mm) along the x-axis, but not along the y-axis (array elevation, 7 mm).

### C. Ultrafast holography of continuous (CW) FUS fields - Experiments in water

The method was first evaluated in degassed water (lossless and homogeneous medium). The density and speed of sound in water at 20°C were respectively assumed to be 1000 kg/m^3^ and 1482.3 m/s [53]. The number of pulse cycles was set to 150, to minimize standing waves.

#### 1. Setup

The experimental setup consisted of an FUS source, and of an ultrasound scanner equipped with a multi-element ultrasound imaging probe (Figure 3). The FUS source was a custom-made single-element spherical concave transducer (PZT, Ferroperm Piezoceramics) with a center frequency of 2.4 MHz, a diameter of 70 mm, and a focal length of 90 mm. The ultrasound receiver was a commercially-available 128-element linear-array imaging probe (ATL L7-4; Philips, Amsterdam, The Netherlands). The footprint of the imaging probe was 38.4 mm × 7 mm (width × elevation), with an element size of 0.3 × 7 mm (pitch × elevation), a frequency bandwidth of 4-7 MHz, and an elevational focus *z*_*f*_ = 25 mm. The source and the receiver array were positioned opposite to each other. The axial distance from the focal position to the sensor surface was set at *z*_*f*_ = 25 mm, so that the elevational focus of the imaging probe and the focus of the source were at the same position (Figure 4a). The sensor array was connected to a programmable ultrafast ultrasound scanner (Vantage 256; Verasonics, Kirkland, WA). This scanner allowed simultaneous acquisition of the incident field on all 128 elements of the sensor array. Acquisition duration ranged 10−100 µs. The source and receiver were synchronized using a trigger signal generated from the scanner. The source was excited by a function generator (AFG3022B; Tektronix, Beaverton, OR) with low amplitude (< 0.7 MPa pressure at the focus) to ensure linear propagation of the incident wave. In every measurement, the acoustic power was below 0.2W.

**Fig. 3.**
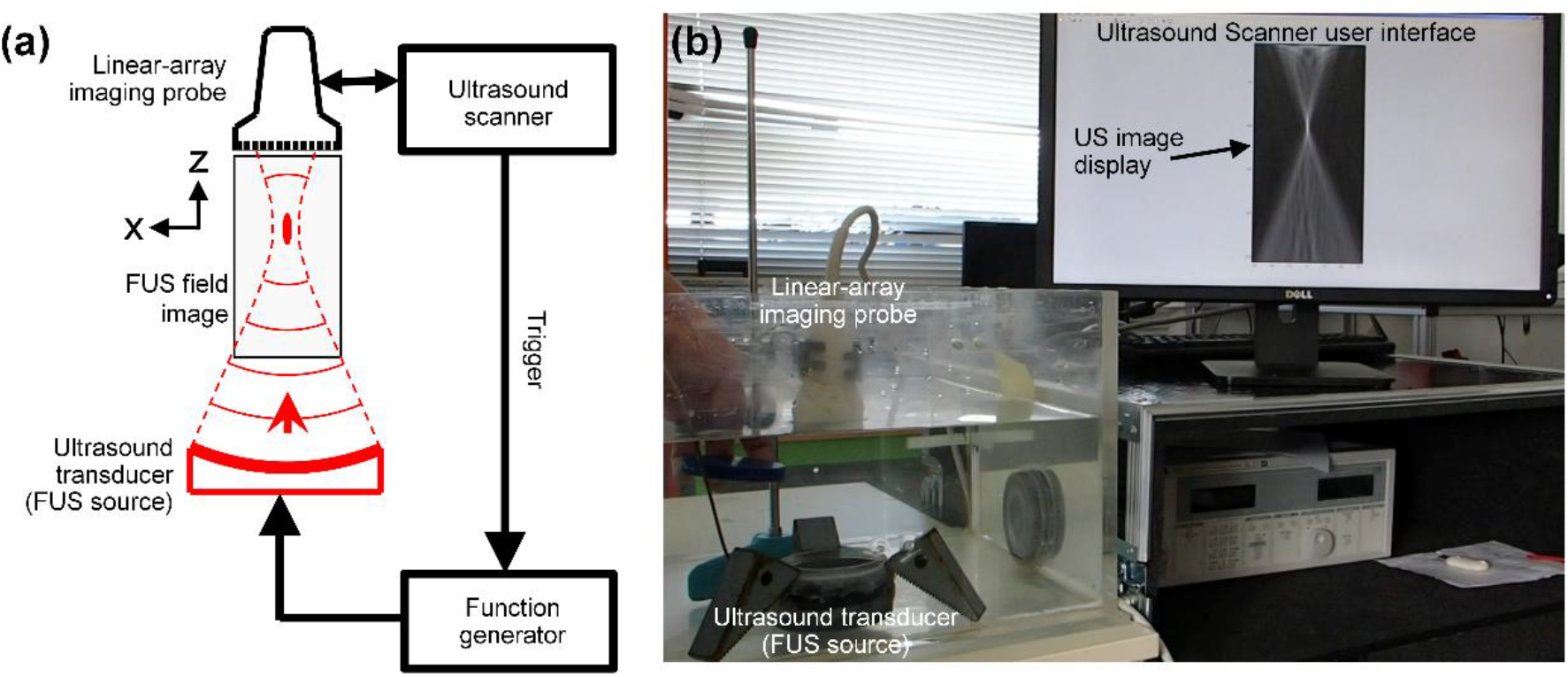
FUS field imaging experimental set-up. An ultrasound imaging array was placed opposite to a FUS transducer, and used as an array of hydrophones, for plane-wave (ultrafast) passive imaging. A portion of the transmitted FUS field was captured simultaneously on all elements of the imaging array. Image series were reconstructed in post-processing using a custom Time Reversal (TR) beamforming. Visualization of the FUS field (envelope) was also feasible in real-time by using the native beamforming of the ultrasound scanner (beamforming for backscattering configuration). **(a)** Set-up schematic. **(b)** Picture of the set-up.

**Fig. 4.**
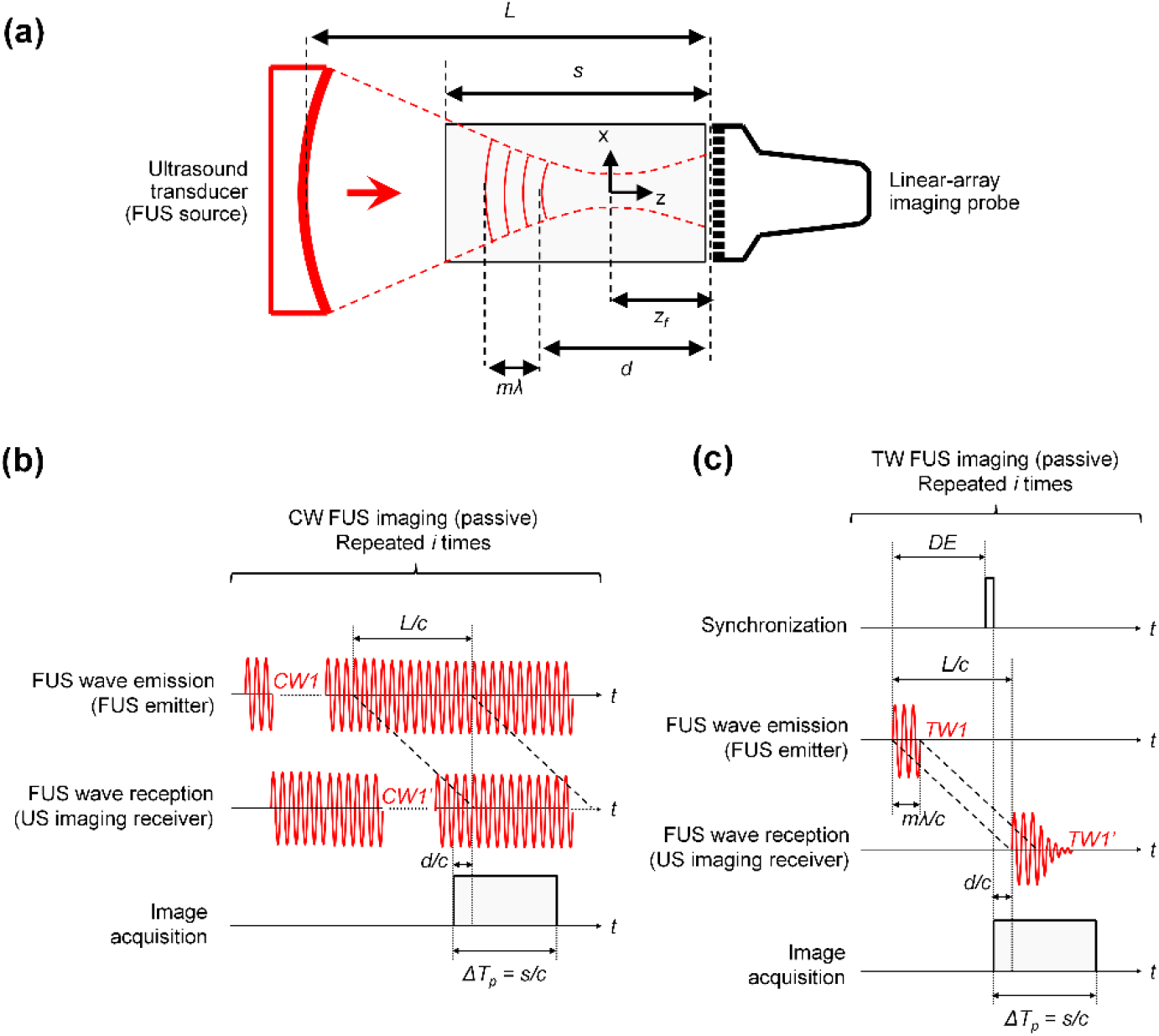
FUS field imaging sequences to visualize ultrasound continuous waves (CW) and/or transient waves (TW) using plane-wave (ultrafast) passive imaging. **(a)** Imaging configuration and associated key parameters for FUS field image reconstruction: *c* the ultrasound celerity; *s* the depth of the FUS field image; d the position of the reconstructed wave front within the FUS field image; R_c_ the radius of curvature / focal length of the FUS emitter; *z*_*f*_ the distance between the FUS focus and the imaging receiver; *L* = *R*_*c*_ + *z*_*f*_ the distance between the FUS emitter and the imaging receiver; *m* the number of ultrasound wavelength (*λ*) per FUS TW emission. **(b)** Chronogram of the FUS emission/reception sequences to visualize CW FUS fields with *CW1* the emitted wave, *CW1’* the received wave and *ΔT*_*p*_ *= s/c* the acquisition duration in passive configuration. No synchronization between the emitter and the receiver was necessary in order to reconstruct an image of the FUS field covering the entire FOV. **(c)** Chronogram of the FUS emission/reception sequences to visualize TW FUS fields with *TW1* the emitted wave, *TW1’* the received wave and *DE* the trigger delay for synchronization of the FUS emitter with the ultrasound scanner. The trigger delay and pulse duration of *TW1* were controlled to reconstruct images of the TW FUS wave event at several instants in time within the FOV (snapshots).

#### 2. Hydrophone measurement (gold standard)

The incident FUS wave field was directly measured by a calibrated needle-like hydrophone (HGL-0200; Onda, Sunnyvale, CA) with a frequency bandwidth of 0.25−40 MHz. Scanning was achieved by a motorized position controller (UMS3; Precision Acoustics, UK) with a spatial step of 0.1 mm and 0.2 mm along the x- and z-axis, respectively.

#### 3. Experimental estimation of the FUS wave field using time reversal

An estimation of the CW fields was reconstructed using the present method of ultrafast holography. The sensor voltage waveforms *u*_*k*_(*t*) were measured experimentally using the imaging probe. In order to minimize standing waves and heating, CW excitations were replaced by long bursts (common practice in conventional hydrophone measurements). The number of pulse cycles of the FUS source was set to 240 cycles (burst duration 100 µs) so that the pulse covers the entire image FOV. Image reconstruction was performed by calculating Equation (8), based on experimental signals (Figure 4b). The image reconstruction was performed offline. The code used for post-processing (Matlab environment) was not optimized for fast reconstruction, contrary to the native beamforming embedded in the ultrasound scanner by the constructor. As a consequence, an additional time of 80 s was necessary for FUS field image reconstruction on an 800 × 1600 pixels grid. The impulse response of the sensor elements *g*_*k*_(*t*) was not known and was arbitrarily set to a Dirac delta function, *g*_*k*_(*t*) = *δ*(*t*), corresponding to 1 m.s^-1^.V^-1^. The reconstructed fields were therefore normalized and reported in arbitrary units (a.u.). The sampling frequency was 20.8 MHz. In Equation (8), cosine interpolation [54] was used to interpolate signal *v*_*k*_ at time *t*+*r*_*k*_/*c*. The spatial sampling interval was 50 µm (∼ λ/12). Note that Equation (8), in the time domain, is clearly not optimized for CW fields. However, an optimized CW monochromatic implementation would miss out on the eventual harmonics that could be present in the received signals. This is why we chose the time domain implementation in this study.

#### 4. Comparison of the estimated CW fields with those expected from simulation B3

Discrepancies between the estimated CW wave field and the hydrophone measurements were expected, because of the suboptimal sensor (linear array). A comparison with simulation B3 is provided, in order to determine how far the experimental results were from the “best achievable result” that can be expected when using this sensor positioned exactly at this position.

### D. Ultrafast holography of transient (TW) FUS fields – Experiments in water

FUS fields in TW mode, commonly used for both ultrasound diagnosis and therapy applications, were also imaged in water.

#### 1. Expected wave field, based on simulations (gold standard)

Hydrophone scanning at all possible positions for all possible time delays was not realistically feasible. Instead of the hydrophone measurements, we provide the expected wave field, based on the impulse response method [52]. The temporal sampling frequency was 500 MHz.

#### 2. Experimental estimation of the FUS wave field using time reversal

Synchronization between the TW FUS emission and the imaging acquisition was necessary to capture the wave moment in the image (Figure 4c). The transient fields were acquired using a single-shot ultrafast acquisition, and reconstructed using Equation (8). Wave movies were created offline, based on a single acquisition, by changing the time t.

#### 3. Comparison of the estimated TW fields with those expected from simulation B3

Discrepancies between the estimated TW wave field and the hydrophone measurements were expected because of the suboptimal setup, too. A simulation was performed in order to determine how far the experimental results were from the “best achievable result” that can be expected when using this sensor positioned exactly at this position. The simulation was essentially the same as simulation B3, except that the continuous wave FUS excitation was replaced by a short burst.

### E. Simultaneous dual-mode experiments in an attenuating tissue mimicking phantom

Next, the feasibility of simultaneous dual mode was tested in a tissue-mimicking gel phantom made of agar and including SiC- and Al2So3 powder [55]. It included cylindrical inclusions (2-cm in diameter) with varying levels of powder. The FUS emitter and imaging array were placed at a distance such that the axial distance between the FUS focal region and the surface of the imaging probe was 25 mm. The phantom was positioned between the FUS emitter and the imaging probe, directly in contact with the latter. For the phantom experiment, a burst duration of 62.5 µs (150 cycles) was used. The density and speed of sound were unchanged assumed to be 1000 kg/m^3^ and 1500 m/s.

The B-mode tissue image was obtained using plane-wave (ultrafast) imaging [56] while the 2.4 MHz FUS source was also emitting. Image registration was automatic since the FUS field and the B-mode images were both acquired simultaneously with the same imaging probe.

The ability to use the resulting real-time image to target the FUS field onto a specific feature, such as an inclusion, was tested, by manually moving the phantom until the FUS focus was at the desired location inside the phantom. A first target location was chosen just at the centre of the inclusion, simulating a localised tumour that needs to be treated. A second location target was chosen far outside of the inclusion, simulating a treatment plan where the inclusion is an organ that needs to be spared.

### F. Simultaneous dual-mode experiments in attenuating soft biological tissues

The same experiment was performed using an ex-vivo porcine loin sample. The burst duration was set to 83.3 µs (200 cycles), and density and speed of sound were respectively assumed to be 1000 kg/m3 and 1540 m/s [57]. No specific target location was defined because no anatomical landmarks could be found in the sample.

### G. Quality metrics

To evaluate the accuracy of the reconstructed FUS field, quality metrics were defined as follows:

- The focal position *FPz* defined as the position of the peak along the z-axis (axial) (Note that the peak positions along the x and y-axis are zero owing to symmetry of the beam),
- The focal length *BWz* defined as the full width at half maximum, along the z-axis, measured at the focal position,
- beam width *BWx* defined as the full width at half maximum, along the x-axis, measured at the focal position

The same quality metrics were used in all simulations and all experiments.

FUS field images were compared to reference FUS fields obtained from gold standard simulations (Rayleigh integral for CW exposures; impulse response for TW exposures) and measurements (raster scan hydrophone). The absolute errors *ΔFPz, ΔBWz*, and *ΔBWx* were calculated. The fraction of FUS energy flux being captured during acquisition was estimated by means of the energy flux ratio *Wr*/*Wi*, which was defined as the ratio of received FUS power *Wr* on the surface of the sensor array to incident FUS power *Wi* measured at the source.

## IV. Results

### A. Approaching the exact FUS field in water in ideal conditions (sim. A1)

Simulation A1 (Figure 5) was performed to assess the feasibility in the ideal case of a 2D-matrix receiving array intercepting all the emitted FUS energy, and with very small elements, in water. The size of the -6 dB focal zone was *BWx* = 1.0 mm in width and *BWz* = 10 mm in length. In these nearly ideal conditions, the energy flux ratio was *Wr*/*Wi* ≈ 99.3%. The simulated FUS field images showed excellent agreement with the reference field (Rayleigh Integral). The absolute errors in the dimensions and position of the -6dB focal zone were Δ*BWx* = 0.4 µm in width, Δ*BWz* = 2.8 µm in length and Δ*FPz* < 25 µm for the focal distance (errors << 0.01%).

**Fig. 5.**
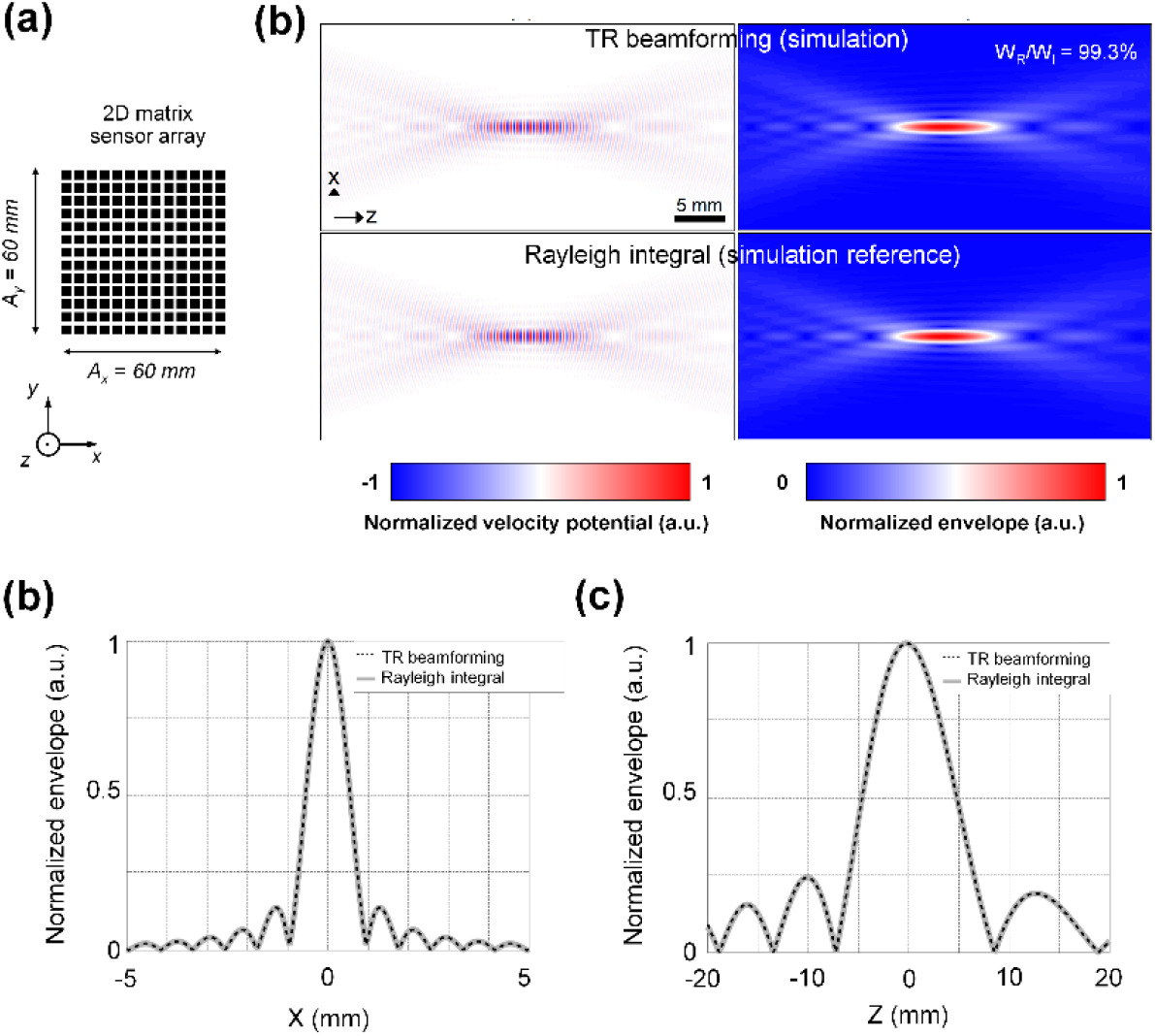
Simulation A1. FUS field imaging in a lossless and homogeneous medium (water) using time reversal (TR) beamforming reconstruction in nearly ideal conditions of FUS energy reception (energy flux ratio *W*_*R*_*/W*_*I*_ *≈ 99*.*3%*) using a 2D-square matrix array receiver (**Table II**). Comparison with the gold standard Rayleigh integral calculation. **(a)** Simulated 2D-array receiver. **(b)** Instantaneous snapshots of the velocity potential and its envelope obtained by TR beamforming (top row) and by solving the Rayleigh integral (reference, bottom row) in the vicinity of the FUS focus. **(c)** Lateral profiles (*x*-axis) of the normalized envelope at *z* = 0 (the focal plane). **(d)** Axial profiles (*z*-axis) of the normalized envelope at *x* = 0 (the focal plane). There was no visible difference between the reference and the simulated TR beamforming method proposed for FUS field imaging.

### B. Holography of FUS fields impacted by suboptimal imaging conditions (Simulations B1-B3)

Simulation B1 (Figure 6) explored the effect of varying the intercepting surface area of the imaging receiver. The reconstruction errors are shown, as a function of the side length of a square sensor array. As expected, the errors in *FPz, BWz*, and *BWx* converged toward zero as the ratio *Wr*/*Wi* tends toward 100%. For this specific position of the sensor array, chosen to be relatively close to the focus, the simulation showed that it is possible to divide the side length by 2 (total surface area divided by 4) without affecting drastically the energy flux ratio (which only decreased from 99.3 to 97.8%). As a consequence, the FUS field could still be reconstructed using a 30×30 mm sensor with negligible errors (Δ*BWx* = 0.4 µm in width, Δ*BWz* = 2.8 µm in length and Δ*FPz* < 25 µm). Below a side length of 30 mm, the absolute errors started to increase significantly (Figure 6a) and to visibly affect the FUS field image (Figure 6b).

**Fig. 6.**
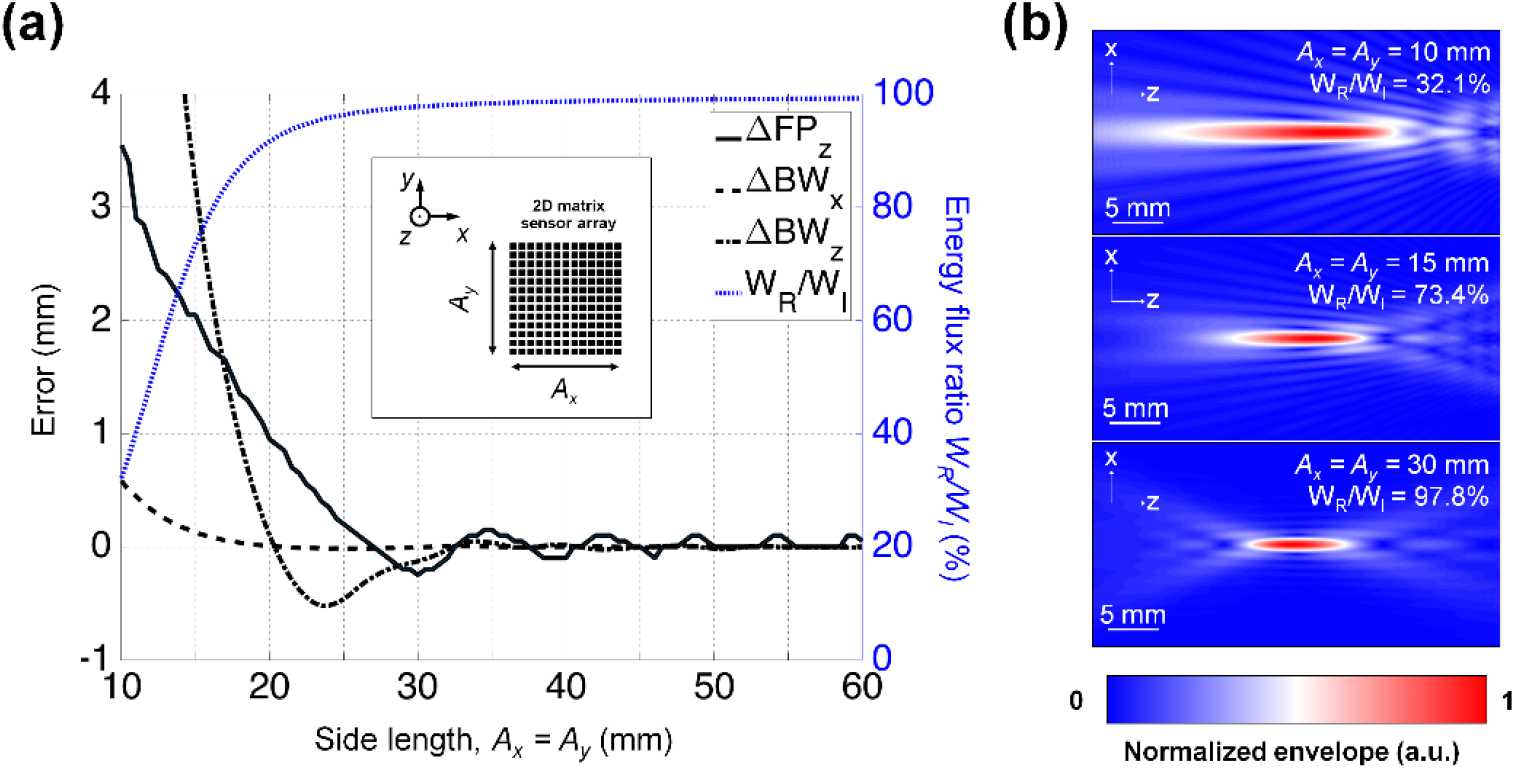
Simulation B1. FUS field imaging in a lossless and homogeneous medium (water) in non-ideal conditions of FUS energy reception (decreasing the energy flux ratio *W*_*R*_*/W*_*I*_) using a 2D-square matrix array with decreasing dimensions in both x and y axis (*A*_*x*_ *= A*_*y*_, **Table II**). **(a)** Influence of the receiver side length on the FUS energy flux ration and on the associated FUS field reconstruction accuracy (reference: Rayleigh integral). **(b)** Envelope of the velocity potential obtained in the vicinity of the FUS focus for 3 cases of imaging receiver dimensions. Reconstruction accuracy improved with increasing energy flux ratio. Note the similarity between the reconstruction obtained for a receiver dimension of *A*_*x*_ *= A*_*y*_ *=* 30 mm and the reference Rayleigh integral calculation (**Fig. 5**)

Simulation B2 (Figure 7) investigated the effect of element size when most of the FUS energy was still captured (99.3%). Reconstruction errors are shown, as a function of the receiver element size. As expected, all error metrics increased with element size increasing from λ/12 to 2.8 λ. As long as the element size was smaller than “λ” /2, the errors in *FPz, BWz*, and *BWx* remained < 0.1 mm, and the FUS field image did not contain visible artifact lobes. For element size between λ/2 and 1.5λ, the absolute errors remained < 0.5 mm, but grating lobes started to be visible. Over 1.5λ, the errors increased significantly with the maxima of the grating lobes getting closer to the focus.

**Fig. 7.**
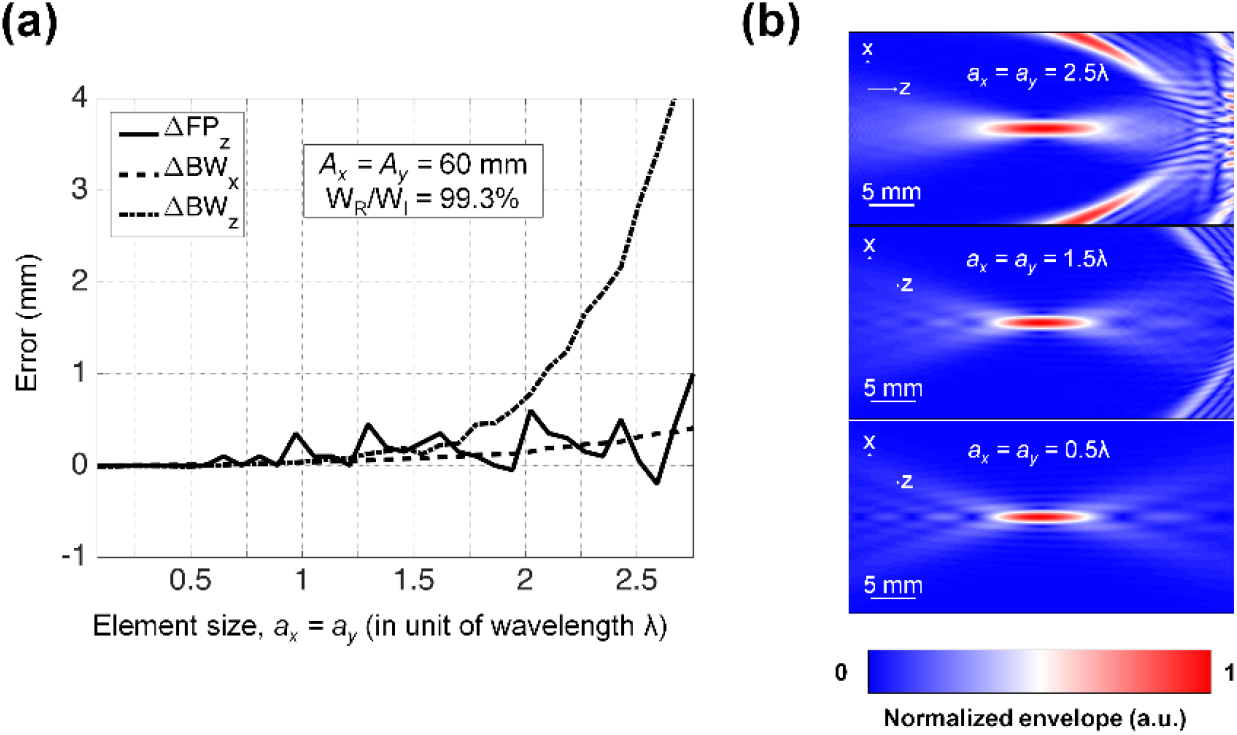
Simulation B2. FUS field imaging in a lossless and homogeneous medium (water) in non-ideal conditions of FUS field acquisition sampling using a 60 mm × 60 mm 2D-matrix array (nearly optimal energy flux ratio of 99.3%) while increasing the elements size *a*_*x*_ *× a*_*y*_ (**Table II**). **(a)** Influence of the receiver element size on the FUS field reconstruction accuracy (reference: Rayleigh integral). **(b)** Envelope of the velocity potential obtained in the vicinity of the FUS focus for 3 cases of imaging receiver element sizes. Images reconstructed with large elements showed artefactual side lobes near the receiver array. Note the similarity between the reconstructed FUS image with element size of 0.5*λ* and the reference Rayleigh integral calculation (**Fig. 5**)

Simulation B3 (Figure 8) investigated the performance of the FUS field imaging when using a standard commercial L7-4 linear-array imaging probe. Such design did not meet the requirements identified in previous sections for achieving optimal FUS imaging. Specifically, the reconstructed FUS fields had larger side lobes compared with the reference fields, due to the low energy flux ratio (*Wr*/*Wi* ≈ 44%) and the large element width along the elevational axis of the sensor array (∼ 11λ). As expected, the images of CW FUS fields showed significant differences in the pre- and post-focal regions, when compared to the reference field (Rayleigh integral). However, the -6 dB focal region was easily identifiable in the image. Its position and dimensions were estimated with an accuracy that might be considered insufficient for an accurate characterization of the device (Δ*BWx* ≈ −0.1 mm in width (10% error), Δ*BWz* ≈ 0.7 mm (7% error) in length and Δ*FPz* ≈ 0.3 mm for the focal distance), but that could be considered sufficient for on-site post-market follow-up of the FUS emitting device, or when using the system for real-time guidance as in subsequent paragraphs.

**Fig. 8.**
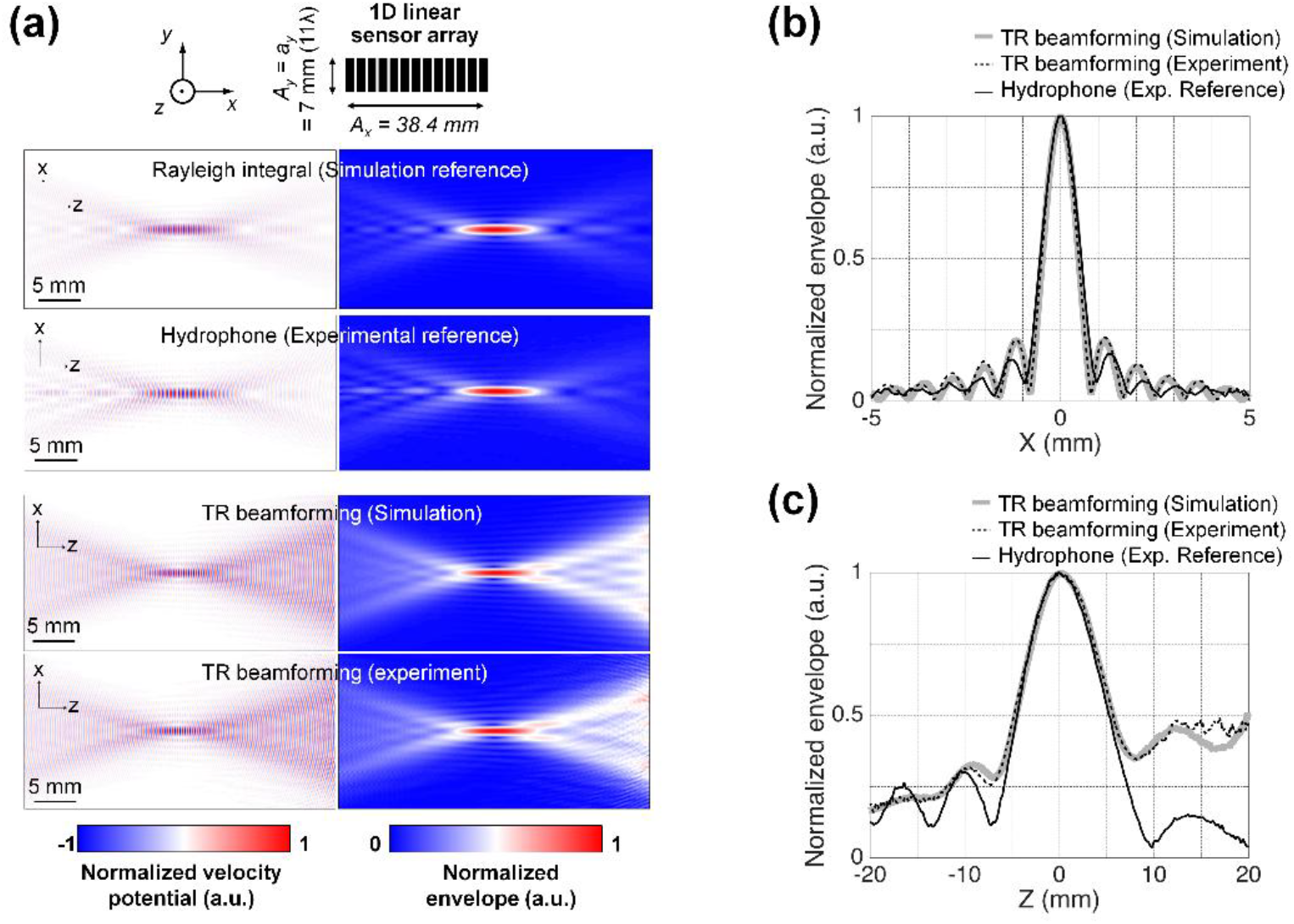
Simulation B3, and corresponding experiment, in water, for a continuous wave (CW), using an ultrasound linear-array probe. **(a)** The instantaneous snapshots of the normalized velocity potential and its envelope obtained in the vicinity of the focus by time reversal (TR) beamforming (Experiment et Simulation, **Scenario #iv, Table II**) wee compared to gold standard: Rayleigh integral (simulation reference) and hydrophone measurement (experimental reference). **(b)** Lateral profiles (*x*-axis) of the normalized envelope at *z* = 0 (the focal plane). **(c)** Axial profiles (*z*-axis) of the normalized envelope at *x* = 0 (the focal plane). Simulated and experimental time reversal data show accurate reconstruction in the focal zone, but imperfect cancellation of the side lobes, especially along the *z*-axis.

### C. Ultrafast holography of CW fields - experimental demonstration in water

The experimental images of the CW FUS field agreed with the prediction (see Figure 8, where simulation B3 and the corresponding experimental measurements are presented side by side). The deviation observed between the experimental image and the true field were therefore expected, and attributable to the suboptimal setup. Despite these differences, the size and position of the focal region itself (main and sides lobes) were still accurately estimated. The absolute errors in the dimensions and position of the -6dB focal zone were Δ*BWx* ≈ −0.1 mm in width (10% error), Δ*BWz* ≈ 0.7 mm (7% error) in length and Δ*FPz* ≈ 0.3 mm for the focal distance. Note that these experimental errors were accurately predicted by simulation B3. Acquisition times for experimental data shown in Figures 8 & 9 ranged 10−100 μs for FUS field imaging, and between 17 min and 28 hours for hydrophone measurements.

### D. Ultrafast holography of TW fields - experimental demonstration in water

A movie showing the propagation of the transient FUS wave was reconstructed (see video in Supplementary material). In this video, the time interval was set to 0.1 µs. The snapshots shown in Figure 9 were extracted from this wave movie, with a coarse interval of 3 µs. Excellent correspondence was observed between the prediction (simulation) and the experimental image. The deviation from the true reference field (i.e., simulation using the impulse response method) was therefore attributable to the suboptimal setup. The acquisition duration for the entire movie was less than 100 µs.

**Fig. 9.**
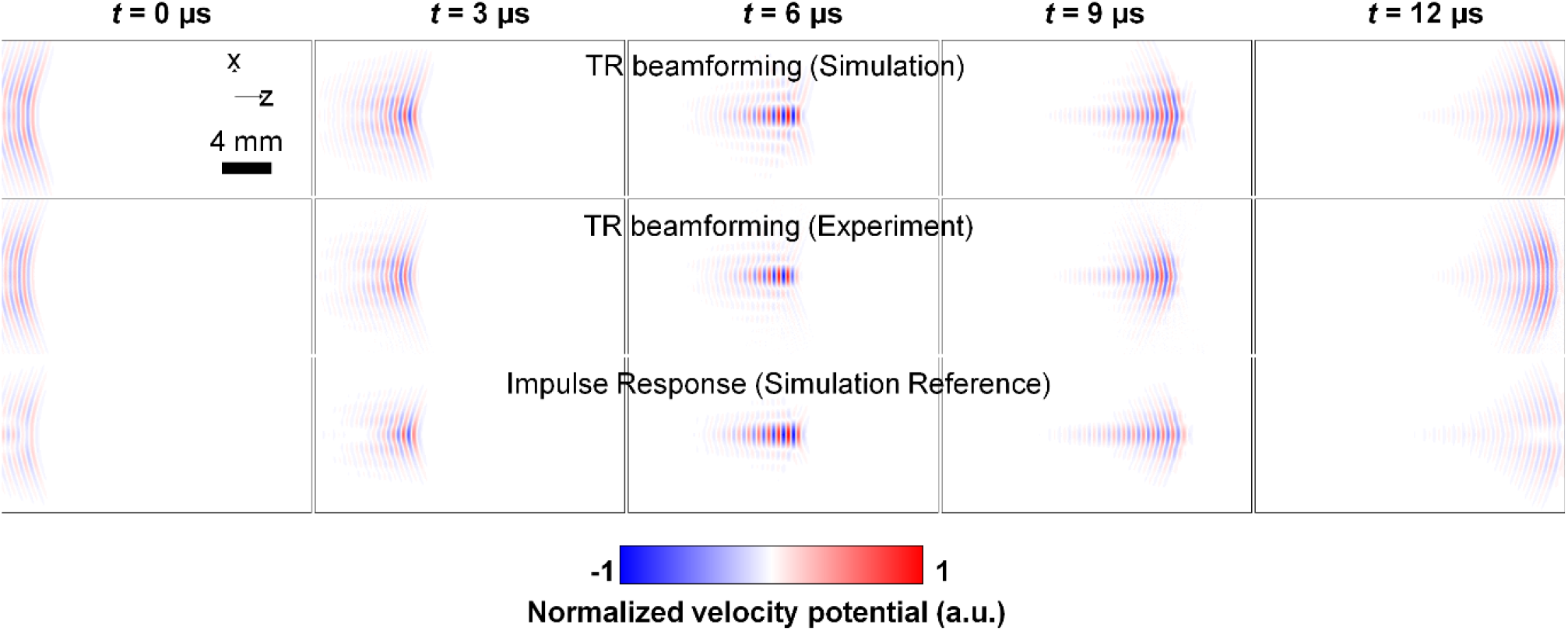
Experiment in water, transient wave (TW). The series of normalized velocity potential in the vicinity of the focus were obtained by time reversal (TR) simulation (top row), TR experiment (middle row), and simulation of the impulse response (simulation reference, bottom row). The temporal interval between images was *dt* = 3 μs. A movie reconstructing the same FUS field propagation with higher temporal sampling (interval *dt* = 0.1 µs) is available (online supplementary material).

### E. Real-time dual mode imaging of FUS fields in a phantom – Experimental demonstration

Experimental simultaneous dual mode images were acquired in a tissue-mimicking gel phantom (Figure 10). These images were obtained and displayed on the scanner in real time, using the built-in pulse echo beamforming algorithm. The wave field displayed in these images is therefore incorrect, because of the use of a suboptimal linear array, but also because of the use of the pulse-echo beamformer (Equation 10) instead of the exact solution (Equation 8), and finally because attenuation was not accounted for. In spite of all these unfavourable conditions, the position of the focal region was easily identifiable. The system was used to target arbitrary regions, whether inside or outside the inclusion. To do so, the phantom was positioned manually. The real-time capability (15 frames per second) of the system made changing the target easy, in just a few seconds.

**Fig. 10.**
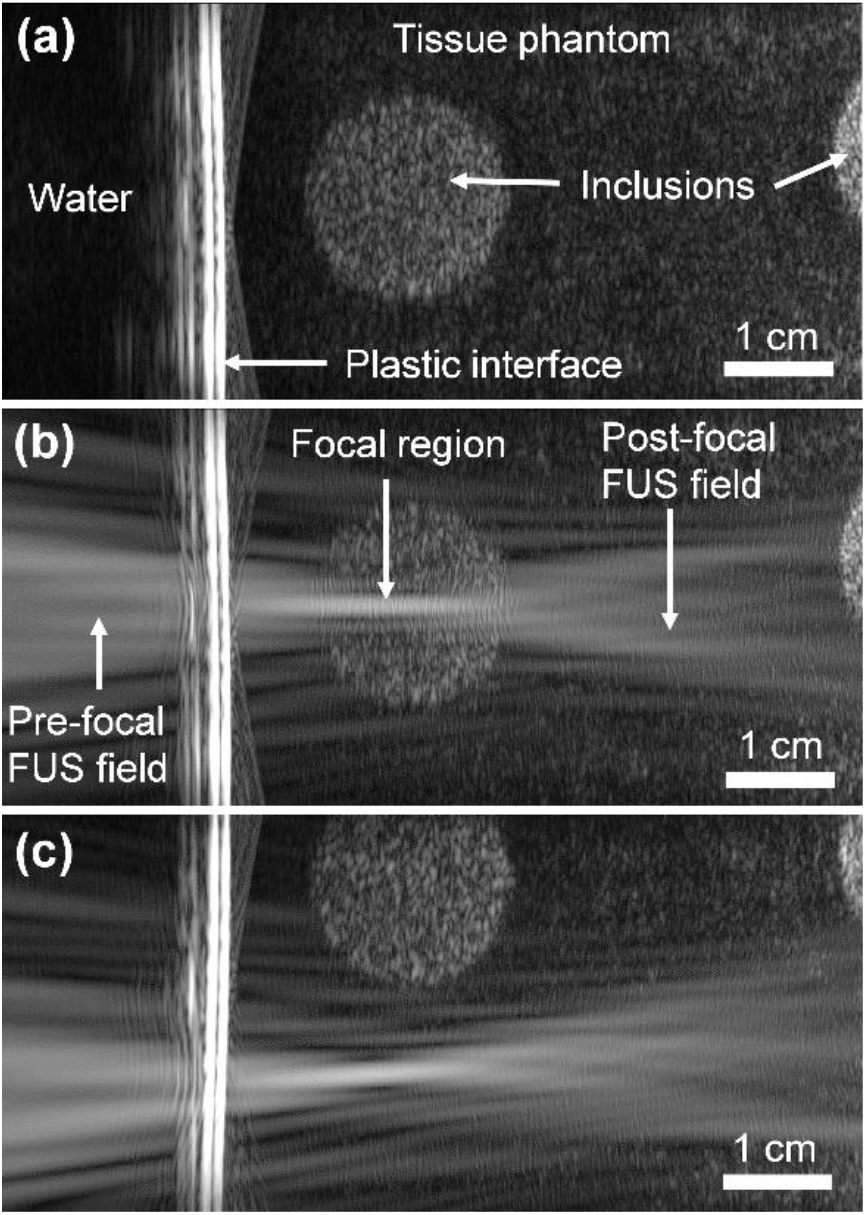
Experiment in a tissue mimicking phantom. *Simultaneous dual mode* imaging allowed real-time positioning of the FUS field relative to a specific target (inclusion). **(a)** Tissue-mimicking gel phantom with a cylindrical hyperechoic inclusion (diameter 20 mm) imaged before FUS wave emission. **(b)** Real-time visualization of a CW FUS field emitted in the phantom. The phantom was positioned manually, using the real-time feedback of the simultaneous dual mode, in order to target the inclusion. **(c)** The phantom was positioned manually, using the real-time feedback, in order to target a region outside the inclusion.

### F. Real-time dual mode imaging of FUS fields in a porcine loin sample – Experimental demonstration

Experimental simultaneous dual mode images were acquired an ex vivo porcine sirloin sample too (Figure 11). These images were also obtained and displayed in real-time. The sample was moved manually in order to target a randomly chosen region of interest. The real-time capability (15 frames per second) of the system made changing the target easy, in just a few seconds.

**Fig. 11.**
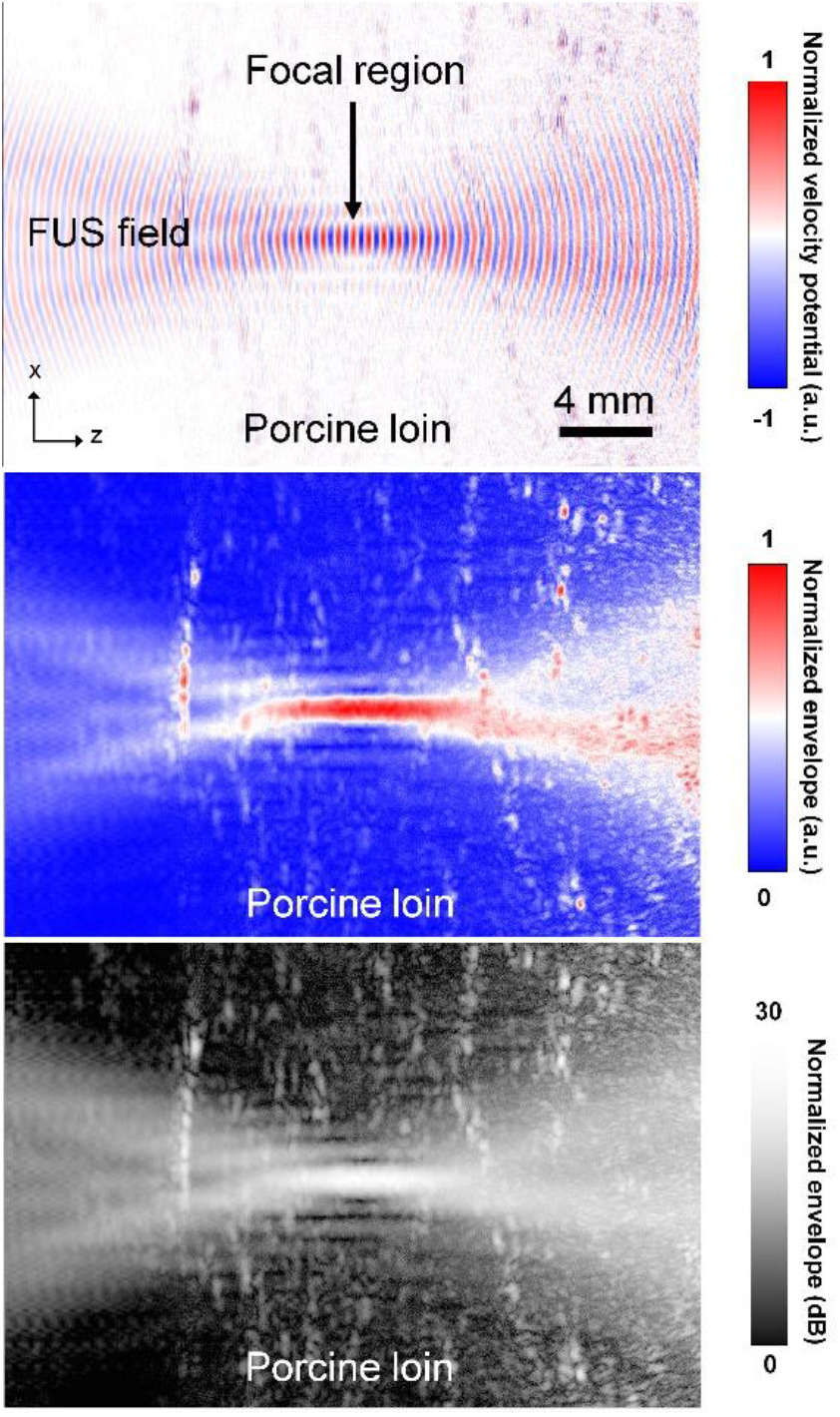
Experiment in a porcine loin sample *ex-vivo*: *dual mode* images using a L7-4 imaging probe, showing a CW FUS field inside a porcine loin sample. The last image (bottom), using *simultaneous dual-mode* acquisition, was available in real-time at 15 frames/sec. It allowed easy guidance to set the FUS focus at any desired position inside the sample. The two other images (top and middle) were obtained independently, using *interleaved dual-mode* acquisition (see Materials & Methods). In these images, the FUS field and the B-mode images are acquired independently. They were also processed offline independently, using Time Reversal for the FUS field, and conventional beamforming for the B-mode. Then the two images were superimposed. Note that the wavelength (approximately 0.6 mm) is consistent with the FUS frequency (2.4 MHz).

## V. Discussion

It is the first time to our knowledge that a relatively accurate visualization of the focal area of a FUS field is provided in real-time at the rates of an ultrasound imaging scanner (Acquisition rate: from 1-1000 frames/s depending of the type of scanner; Display rate: 10-15 frames/s). The FUS field was visible using a simple 1D linear array, affordable (not more expansive than hydrophones) and available in many research laboratories and hospitals. The 1D-linear imaging array used experimentally was affordable, but clearly suboptimal. Specifically, energy flux ratio was < 50%, and element size along the elevational axis was >> λ/2. Despite these unfavourable conditions, real-time ultrafast holography of FUS fields method remained feasible and informative in water and in biological tissues. Especially, the FUS beam width and focal length, two main features of the FUS field that need to be monitored during medical procedures, were accurately estimated.

Two cases were investigated with transient FUS waves typically used for ultrasound imaging, or for non-thermal ultrasound therapies (e.g., lithotripsy, histotripsy, BBB opening, neurostimulation), and continuous FUS waves typically used for thermal ultrasound therapies (e.g. high intensity focused ultrasound - HIFU).

The main limitation observed with such a suboptimal receiver array was that quantitative estimation of the pressure level was not possible. But even in these suboptimal conditions, the real-time nature of the technique makes it fit for some applications, such as on-site quality control and follow up of devices that are installed in the hospital.

In the future, better and quantitative images can be obtained with the development of specific matrix arrays, or by moving a linear array composed of small elements.

The first potential application is fast ultrasound metrology, replacing the lengthy hydrophone scans. In the past, holography has been used to detect the eventual flaws, by imaging the vibration at the surface of the source, at the expense of lengthy scans [14]. We anticipate that this application can now be made real time.

Finally, in some very specific therapeutic or industrial applications where it is possible to position the imaging array opposite to the FUS transducer, the technique may also provide real-time feedback and targeting capability. The use of a thin acoustic mirror, or of thin flexible sensors, may also enable FUS field imaging in applications where it is not possible to position an imaging probe opposition to the FUS transducer.

Just as a note, in our laboratory, real time FUS field imaging was also implemented on a conventional low-speed ultrasound scanner that scans the field of view line-by-line. However, reporting on this implementation is beyond the scope of this article.

Finally, the method is not appropriate for diverging wave fields. Such fields would require a full enclosure with small elements. The complexity and cost of such a device would be prohibitive.

## VI. Conclusion

This paper presents a method and technical proof-of-concept for real-time imaging of focused ultrasound (FUS) fields, in water and in soft biological tissues. Accurate instantaneous snapshot of FUS fields were reconstructed, using an ultrafast ultrasound imaging system. “FUS wave movies” showing the propagation of the wave were also presented. The prerequisite and feasibility of the method were investigated in a favourable condition (water) while preliminary trials were initiated in soft biological tissues. The presented method provides a novel procedure to characterize FUS fields, qualitatively in most conditions, but with potential for quantitative measurements in water. This technique is also a novel opportunity to guide in real time FUS fields in biological soft tissues, with potential interest for the guidance of certain therapeutic ultrasound devices and procedures.

## Supporting information

Movie showing the transient FUS wave

## Acknowledgements

The authors thank Dr. Katrin Skerl for preparing the phantoms. Rémi Souchon warmly dedicates this work to his beloved son Armand, who was present at LabTAU during the initial works in 2016, and whose passing away during these days contributed to delaying its publication so much.

## Appendix A. The pulse-echo approximation

The pulse-echo (conventional beamformer) approximation is of special interest because it is readily available in real time in all medical ultrasound scanners. It is therefore worth investigating how accurate that approximation is. In this appendix, we derive the relation between conventional beamforming (10) and Time Reversal (8), and highlight how the conventional beamforming can approximate the wave field of interest. For the sake of convenience, these two equations are recalled below:

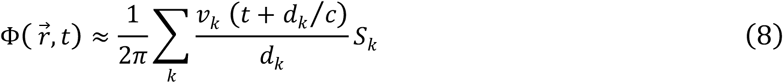

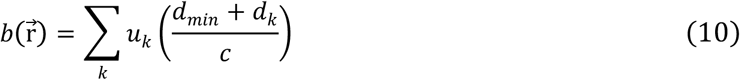

### A. Echo Ranging

In equations (8) and (10), the point of observation can be chosen arbitrarily. However, depending on whether we use a conventional beamformer or Time Reversal, the associated time delays will be different.

Let vector 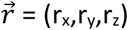 be the point of observation (we choose it). The distance *d*_*min*_ corresponds to the shortest path between the imaging probe and the point of observation. It determines the time of arrival of the acoustic wave at that point: *t*_*min*_ = *d*_*min*_ /*c*, with *c* the speed of sound. The distance *d*_*k*_ corresponds to the path between the point of observation and the element number *k*. It determines the delay of arrival of the backscattered wave on the *k*-th element: *t*_*back*_ = *d*_*k*_ /*c*. The total travel path is *d*_*min*_+*d*_*k*_, and the corresponding delay is (*d*_*min*_+*d*_*k*_)/*c*, yielding the standard beamforming equation (10).

In a conventional beamformer, the earliest associated time delay will be *τ*_*b*_ = 2*d*_*min*_/*c*, and this equation is used to match the corresponding position in space. For a given delay *τ*_*b*_, the associated position will be at depth *d*_*min*_ = *cτ*_*b*_ /2. However, in the case of Time Reversal, the associated time delay is simply *τ*_*TR*_ = *d*_*k*_/*c*, the associated position will be at depth *d*_*min*_ = *cτ*_*TR*_. The consequence of using the conventional beamforming equation (10) on wave signals arising from an external source will therefore be a spatial scaling, similar to a compression by a factor of two, in the direction normal to the imaging probe. For example, a transient burst emitted from a source located 10 cm away from the imaging probe will be displayed at a depth of 5 cm. The wavelength will also be compressed by a factor of two.

However, and surprisingly, our experience with simultaneous dual-mode showed that spatial variations of the wave field, such as the position of the focus and the shape of the wave field, were unaffected. In order to understand why, it is necessary to study the delay laws.

### B. The delay laws

In a conventional beamformer, the time of flight is equal to *t*_*conv*_ = (*d*_*min*_+*d*_*k*_)/*c*. The shortest time of flight is along the path that returns to the nearest element, namely 2*d*_*min*_/*c*. The corresponding delay law, i.e. the delays to be applied to the individual elements, is given by the difference: *τ*_*conv*_ = *t*_*conv*_ - 2*d*_*min*_/*c =* (*d*_*k*_-*d*_*min*_)/*c*.

In the Time Reversal equation, the time of flight is equal to *t*_*TR*_ = *d*_*k*_/*c*. The shortest time of flight is along the path that returns to the nearest element, namely *d*_*min*_/*c*. The corresponding delay law, i.e. the delays to be applied to the individual elements, is given by the difference: *τ*_*TR*_ = *t*_*TR*_ - *d*_*min*_/*c =* (*d*_*k*_-*d*_*min*_)/*c*.

It turns out that the delay law is the same, in both equations. This is why the conventional B-mode image displays the focus of the HIFU beam at the correct position, even though the equation (10) should not be used.

This equality is illustrated in a simple example below (Table III). The numbers are drawn from the L7-4 imaging probe used throughout the article. Its pitch is 0.3 mm, and speed of sound was assumed to be 1500 m/s. For an observation point placed at depth z=25 m. The time of flight and the delay law was calculated for some of the elements, for conventional beamforming (a), and for Time Reversal (b).

### C. The beamforming equations

Let us now introduce three simplifying assumptions:

a. All elements have the same sensitivity, and that sensitivity is constant over the frequency bandwidth
b. All elements have the same surface.
c. The point of observation is located sufficiently far away from the imaging probe.

Based on Assumption (a), and using the plane wave assumption (*p*=*ρcv*), we can write:

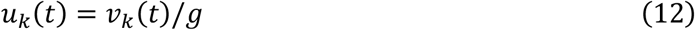

where *v*_*k*_(*t*) is the normal velocity on the *k*-th element, *g* is the sensitivity of the elements (expressed in V per m.s^-1^). For hydrophones, *g* can be calculated from the sensitivity data, commonly given in units of Pa/V, and using the plane wave equation *p*=*ρcv*. For imaging probes, this value is generally not communicated, but it can be measured exactly the same way as we would for a hydrophone.

Based on Assumption (b), and noting S the total surface of the imaging probe, we can write:

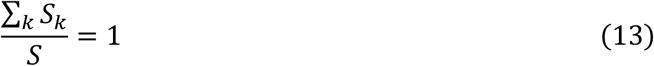

Now based on Assumption (c), let *ε*_*k*_ = *d*_*k*_ – *d*_*min*_ be the difference between the forward path and the backpropagating path. This difference can be significant near the imaging probe (i.e. when *d*_*min*_ tends towards zero). The difference becomes smaller and smaller as the distance *d*_*min*_ increases and the point of observation moves away from the imaging probe. For distances *d*_*min*_ greater than the effective aperture of the imaging probe, the difference *ε*_*k*_ becomes negligible. Then, for any value of *k*:

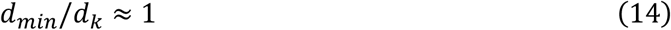

Plugging consecutively (12), (13), and (14) into equation (10) yields:

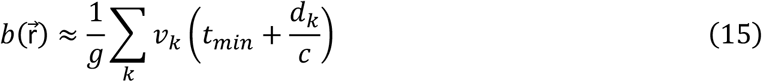

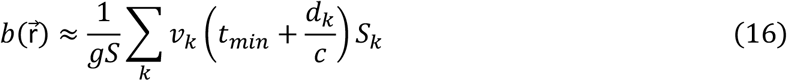

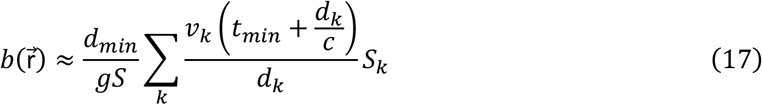

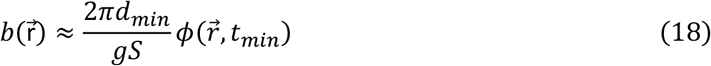

In plain words, the conventional beamformed image 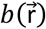 is an approximation of the wave field ϕ, observed at time *t* = *t*_min_. However, it should be recalled the depth *d*_min_ and the corresponding time *t*_min_ actually change with the point of observation. Strictly speaking, they should be noted 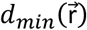 and 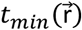 respectively:

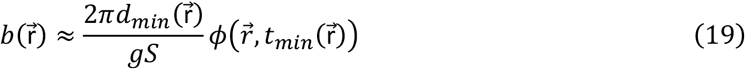

As depth increases, the delay *t*_min_ also increases. The deeper we observe, the “later in time” we see the wave. The result is somewhat similar in essence to the red shift in spatial observations, where looking at further distance implies looking back in time, at older phenomena. Except that the phenomenon is now in the opposite direction: looking further away implies looking into the future of the wave. This behaviour should actually be no surprise to ultrasound physicists, who know that deeper echoes are actually generated at a later time. The main consequence is that the conventional B-mode image is not an “instantaneous snapshot” of the wave field at some fixed time.

### D. Hacking TGC

Two interesting consequences can be inferred from equation (19). First, the amplitude of the wave field can be recovered by compensating its dependence on depth. On a conventional ultrasound scanner, it can be done by setting the built-in Time Gain Compensation (TGC) settings to match and counter-balance that dependence, with:

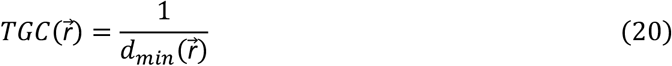

The TGC-compensated B-mode image will now be:

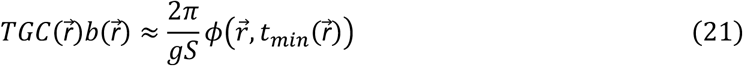

Second, for continuous wave excitation, it can be noted that the use of the incorrect time delay only affects the phase of the signal. Its envelope, i.e. the amplitude of the field, is unchanged:

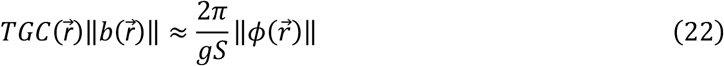

Equation (22) has practical applications, because the envelope 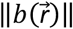 of the beamformed signals is the entity being displayed by conventional ultrasound scanners. The amplitude of the HIFU field can therefore be observed directly, provided that the TGC is set appropriately.

