## Supplementary material for "Real-time ultrafast holography of focused ultrasound wave fields using an ultrasound scanner": Tables

| **Symbol** | **Description** |
| --- | --- |
|  | OPERATORS |
| $*$ | Convolution operator |
| $\text{∇}$ | Gradient operator |
| $\text{∂}$ | Partial derivative operator |
|  | SCALAR VALUES |
| *c* | Speed of sound |
| *d* | *d* $\text{=}\text{ }\left\Vert\vec{\text{r}} \text{-}\text{ }\vec{\text{r}_{\text{s}}} \right\Vert$ is the distance between the elementary integration surface $\vec{dS}$ and the observation position $\vec{\text{r}}$ |
| $d_{\text{k}}$ | Distance between the observation position $\vec{\text{r}}$ and the center of the *k*-th element of the sensor array |
| $\text{e}_{\text{k}}\text{(}\text{t}\text{)}$ | Electrical voltage applied to the *k*-th element when working in transmit mode |
| $\text{g}_{\text{k}}\text{(}\text{t}\text{)}$ | Acousto-electric impulse response on the *k*-th element of the sensor array, when working in receive mode, and defined in units of m.s^-1^.V^-1^ |
| $\text{h}_{\text{k}}\text{(}\text{t}\text{)}$ | Electro-acoustic impulse response on the *k*-th element of the sensor array, when working in transmit mode, and defined in units of V.m^-1^.s |
| *p* | Pressure |
| *S* | Total surface of the sensor array |
| $\text{S}_{\text{k}}$ | Surface of the *k*-th element of the sensor array |
| *t* | Time |
| *T* | Total duration of the experiment |
| $\text{u}_{\text{k}}\text{(}\text{t}\text{)}$ | Electrical voltage measured on the *k*-th element when working in receive mode |
| $\text{v}_{\text{k}}\text{(}\text{t}\text{)}$ | Normal particle velocity measured on the *k*-th element |
| $\text{v}_{\text{k}}^{\text{TR}}\left( \vec{\text{r}_{\text{s}}}\text{,}\text{ }\text{t} \right)$ | Time-reversed normal velocity on the surface of the sensor array, with $\text{v}^{\text{TR}}\left( \vec{\text{r}_{\text{s}}}\text{, }\text{t} \right)=\vec{v}^{TR}\left( \vec{\text{r}_{\text{s}}}\text{, }\text{t} \right)\cdot\vec{n}_{k}$ |
| $\text{Φ}\left( \vec{\text{r}}\text{, }\text{t} \right)$ | Velocity potential of the incident field at observation position $\vec{\text{r}}$ and time *t* |
| $\text{Φ}^{\text{TR}}\left( \vec{\text{r}}\text{,}\text{ }\text{t} \right)$ | Velocity potential of the time-reversed field at observation position $\vec{\text{r}}$ and time *t* |
| *ρ* | Density |
|  | VECTORS |
| $\vec{dS}$ | Elementary integration surface, on the sensor array, with $\vec{dS}=dS.\vec{n}$ |
| $\vec{\text{I}}$ | Acoustic intensity (Poynting vector) |
| $\vec{\text{n}}$ | Unit vector, normal to the elementary integration surface *dS*, and pointing from the integration surface toward the medium |
| $\vec{n}_{k}$ | Unit vector, normal to the k-th surface element, and pointing from the sensor toward the medium |
| $\vec{\text{r}}$ | Observation position |
| $\vec{r}_{S}$ | Position of the elementary integration surface $\vec{dS}$ on the sensor array |
| $\vec{\text{v}}$ | Particle velocity |
| $\vec{v}^{TR}\left( \vec{\text{r}_{\text{s}}}\text{,}\text{ }\text{t} \right)$ | Time-reversed particle velocity vector on the surface of the sensor array |
| $\vec{v}_{\text{k}}^{\text{TR}}(t)$ | Time-reversed normal particle velocity applied to the *k*-th element |

Table I. Symbols, grouped by type (operators, scalar, vectors), and sorted by alphabetical order

| **Simulation #** | **A1. Optimal matrix** | **B1. Suboptimal surface** | **B2. Suboptimal element size** | **B3. Linear array** |
| --- | --- | --- | --- | --- |
| Type of imaging array | 2D Matrix | 2D Matrix | 2D Matrix | 1D Linear |
| Imaging array dimensions, *A_x_* × *A_y_* (mm × mm) | 60×60 | Variable  10×10 – 60×60 | 60×60 | Rectangular,  38.4×7 |
| Element size, *a_x_* × *a_y_*  (mm × mm)  [wavelengths] | Square,  0.05×0.05 [*λ*/12×*λ*/12] | Square, 0.05×0.05 [*λ*/12×*λ*/12] | Square, Variable  0.05×0.05 – 1.7×1.7  [*λ*/12×*λ*/12 – 2.8*λ*×2.8*λ*] | Rectangular,  0.3×7  [*λ*/2×11*λ*] |

Table II. FUS field imaging configurations investigated in the numerical simulations

| Element | -4 | -3 | -2 | 1 | 0 | 1 | 2 | 3 | 4 |
| --- | --- | --- | --- | --- | --- | --- | --- | --- | --- |
| **Conventional beamforming** | | | | | | | | | |
| Time of flight (µs) | 33.3525 | 33.3441 | 33.3381 | 33.3345 | 33.3333 | 33.3345 | 33.3381 | 33.3441 | 33.3525 |
| Delay law (ns) | 19.1890 | 10.7965 | 4.7993 | 1.2000 | 0 | 1.2000 | 4.7993 | 10.7965 | 19.1890 |
| **Time Reversal** | | | | | | | | | |
| Time of flight (µs) | 16.6859 | 16.6775 | 16.6715 | 16.6679 | 16.6667 | 16.6679 | 16.6715 | 16.6775 | 16.6859 |
| Delay law (ns) | 19.1890 | 10.7965 | 4.7993 | 1.2000 | 0 | 1.2000 | 4.7993 | 10.7965 | 19.1890 |

Table III. Time of flight, and delay law, for a point of observation located at depth z=25 mm from the imaging probe. Note how the delay law is the same for conventional beamforming and for Time Reversal, even though the times of flight are different.
